# Eurasian Stone-curlews Burhinus oedicnemus breeding in Italy show a remarkable inter and intra-population variability of migratory behaviour

**DOI:** 10.1101/2022.02.17.480878

**Authors:** Valentina Falchi, Andrea Barbon, Carlo Catoni, Giulia Cerritelli, Alice Cutroneo, Giacomo Dell’omo, Marco Dragonetti, Pietro Giovacchini, Angelo Meschini, Lucio Panzarin, Angela Picciau, Dimitri Giunchi

## Abstract

Migratory behaviour in birds shows a remarkable variability at species, population and individual levels. Short-distance migrants, in particular, often adopt a partial migratory strategy and tend to have a rather flexible migration schedule which allows them to respond more effectively to extreme environmental variations, like those due to climate change. Weather seasonality and environmental heterogeneity at regional and local scales have been reported as significant factors in the diversification of migratory behaviour for some species of Mediterranean migrants. Relatively few studies, however, investigated the migration patterns of non-passerine birds migrating within this area. In this study we investigated the migratory strategy of the Eurasian Stone-curlew *Burhinus oedicnemus* using data collected on 40 individuals tagged with geolocators and GPS-GSM tags, belonging to two continental and two Mediterranean populations of the Italian peninsula. The proportion of migrants was significantly higher in continental populations, but we observed a significant variability also within Mediterranean populations. GPS-tagged migrants traveled less than 1000 km, spending the winter within the Mediterranean basin. Continental Stone-curlews i) departed earlier in spring and later in autumn and ii) covered longer distances than those from Mediterranean areas. The speed of migration did not change between seasons for continental birds, while Mediterranean individuals tended to migrate faster in spring. The likelihood of departure for autumn migration of GPS-tagged birds significantly increased when temperatures were near or below 0 °C suggesting that Stone-curlews tend to delay departure weather conditions worsen abruptly. Thus it can be speculated that the frequency of migratory birds in the considered populations may decrease in the near future due to the effect of global warming in the Mediterranean. This could have a significant effect on the distribution of species throughout the year and should be taken into account when targeting conservation measures.

## Introduction

The extent and the pattern of migration may vary strongly among different taxa and even within the same taxon (Dingle & Drake 2007). Migratory behaviour in birds, in particular, shows a remarkable variability at species (Schmaljohann 2018, Anderson *et al*. 2020), population (Monti *et al*. 2018, Phipps *et al*. 2019) and individual levels (Shamoun-Baranes *et al*. 2017, Tedeschi *et al*. 2019). One of the most extreme forms of intra-population variation in migratory behaviour is partial migration, where only a fraction of individuals are migrants (Chapman *et al*. 2011, 2014). This seems to be the most common strategy in birds (Chapman *et al*. 2011), especially among short-distance migrants (Pulido *et al*. 1996, Newton 2008).

Besides intra-population heterogeneity in migratory strategy, short-distance migrants also show considerable variability in migratory behaviour, mainly due to the effect of environmental factors in modulating the phenology of migration (Berthold 1996, Pulido & Widmer 2005, Both *et al*. 2010, Newton 2012, Åkesson *et al*. 2017). This is especially true for post-breeding migration as birds do not have the urgency to get to breeding grounds first and in time (Kokko 1999, Nilsson *et al*. 2013). Due to the relative proximity between breeding and wintering grounds, short-distance migrants can also optimize their departure schedules by selecting weather conditions that allow them to minimize energy expenditure throughout the trip (Alerstam & Lindstrom 1990, Lehikoinen *et al*. 2004, Rubolini *et al*. 2007, Møller *et al*. 2010, Knudsen *et al*. 2011, Nilsson *et al*. 2013). This leads to more flexible migration schedules for short-distance than for long-distance migrants (Pulido & Widmer 2005, Dingemanse & Wolf 2013, Åkesson *et al*. 2017) and allows the former to respond more effectively to extreme environmental variations, like those due to climate change (Jenni & Kéry 2003, Gordo 2007).

Weather seasonality and environmental heterogeneity at regional and local scales are significant factors in the diversification of migratory behaviour at inter and intra-population levels for some species migrating within the Mediterranean region (de La Morena *et al*. 2015, Monti *et al*. 2018). Moreover, the ongoing increase in temperature recorded in this area due to global warming (Mariotti *et al*. 2015) can affect migration timing of short distance migrants (e.g. Birtsas *et al*. 2013) and might even lead to the complete suppression of their migratory behaviour (Jenni & Kéry 2003, Gordo 2007). Relatively few studies, however, have investigated migration patterns and the role of environmental factors in modulating migratory behaviour of non-passerine birds migrating within this area (but see Birtsas *et al*. 2013, de La Morena *et al*. 2015, Monti *et al*. 2018; Alonso *et al*. 2019).

The Eurasian Stone-curlew *Burhinus oedicnemus* (Charadriiformes, Burhinidae, hereafter, Stonecurlew) is a steppic bird whose European populations are mainly concentrated in the Mediterranean area (Vaughan & Vaughan-Jennings 2005, Delany *et al*. 2009, Keller *et al*. 2020). In Italy the species is relatively widespread in Sardinia, Sicily and in the Southern part of the peninsula, while it is scarce, although locally common, along the riverbanks and in farmlands in the Centre and the North of the peninsula (Brichetti & Fracasso 2004, Biondi & Pietrelli 2015).

The species is of European conservation concern (BirdLife International 2021) but information on the status of its populations is limited (Gaget *et al*. 2019). Available information suggests that the migratory behaviour of the Stone-curlew should be rather variable at intra- and inter-population level throughout its range (Cramp & Simmons 1983, Green *et al*. 1997, Vaughan & Vaughan-Jennings 2005, Giunchi *et al*. 2015), but relatively little is known about its movements. Investigating the migratory strategies of different populations, the geographical range of their movements and the degree of migratory connectivity is an important contribution for understanding the factors affecting populations trends (Webster *et al*. 2002, Taylor & Norris 2010, Finch *et al*. 2017). Furthermore, understanding how migration phenology is influenced by proximate meteorological factors can help to understand how a species would react to the ongoing climate change (Gill *et al*. 2014, Haest *et al*. 2018, Burnside *et al*. 2021, Linek *et al*. 2021).

In this study we 1) investigated the spring and autumn migratory strategies of Stone-curlews belonging to four Italian populations, two from the North and two from the center of the peninsula; 2) performed a preliminary evaluation of the possible role of exogenous factors in modulating the phenology of autumn migration. We expected a higher proportion of migrants in northern populations than in southern ones, as reported for other Palearctic migrants (Newton & Dale 1996, Newton 2008, Linek *et al*. 2021) and suggested for the Stone-curlew (Delany *et al*. 2009, Biondi & Pietrelli 2015). Northern breeding birds should leave their wintering/breeding areas earlier than birds breeding more in the South as observed in other Palearctic species (Newton & Dale 1996, Newton 2008, Ambrosini *et al*. 2016, Briedis *et al*. 2020). Stone-curlews should use a timeminimization strategy during spring migration and an energy-minimization strategy during autumn migration (Alerstam & Lindström 1990, Kokko 1999, Nilsson *et al*. 2013) and thus spring migration is expected to be faster than autumn migration in all populations, as reported for several bird species, including waders (Zhao *et al*. 2017, Schmaljohann 2018). Breeding latitude should affect the timing of departure dates, i.e. birds from northern populations are expected to depart earlier in autumn and later in spring with respect to birds from southern populations (Newton 2008, Conklin *et al*. 2010, Linek *et al*. 2021). The timing of migration was expected to be quite variable,as short-distance migrants usually show a quite flexible migratory schedule (Pulido & Widmer 2005, Newton 2008, Pulido & Berthold 2010, Monti *et al*. 2018b); this variability should be particularly high in autumn, when there is no compelling need to reach the wintering sites (Shamoun-Baranes *et al*. 2006, Nilsson *et al*. 2013). Autumn migration was expected to be significantly affected by meteorological variables, such as wind conditions (Liechti 2006, Conklin & Battley 2011, Grönroos *et al*. 2012, Duijns *et al*. 2017), temperature (Burnside *at al*. 2021), and atmospheric pressure (Dänhardt & Lindström 2001). Low temperatures, in particular, should significantly increase the likelihood of migratory departure, given 1) their negative effect on the availability of invertebrates (Eggleton *et al*. 2009, Abram *et al.,* 2017) which are the main target preys for the Stone-curlew (Amat 1986, Green *et al*. 2000, Giannangeli *et al*. 2004, Giovacchini *et al*. 2017), and 2) the low resting metabolic rate of the Stone-curlew (Duriez *et al*. 2010), which could make the thermoregulation process in cold conditions rather costly.

## Methods

### Study areas

The study was carried out on four Italian areas, two located in the Continental biogeographical region (https://www.eea.europa.eu/data-and-maps/data/biogeographical-regions-europe-3), Taro river (Parma Province, Northern Italy, 44.74°N 10.17°E; n = 9) and Piave river (Treviso Province, Northern Italy, 45.80°N 12.28°E; n = 12), and two in the Mediterranean biogeographical region, Grosseto Province (Southern Tuscany, Central Italy, 42.60°N 11.22°E; n = 10) and Viterbo Province (Northern Latium, Central Italy, 42.35°N 11.83°E; n = 10) (Fig. 1). According to the Köppen-Geiger climate classification, which is based on the relationship between climate and vegetation (Köppen 2011, Chen & Chen 2013), Taro and Piave areas are within a mild temperate, fully humid zone with warm summer (warmest month average temperature <22 °C and at least four months with temperatures ≥10 °C), while Grosseto and Viterbo areas are within a mild temperate zone with dry and hot summer (warmest month average temperature ≥22 °C) (data obtained using the R-package ‘kgc’ 1.0.02, Bryant *et al*. 2017). Birds were captured during the breeding period (March-July) from 2013 to 2019 using different methods (i.e. mist nets, fall traps, dip nets). All birds were measured according to standard ringing procedures (Busse & Meissner 2015) and molecularly sexed according to Griffiths *et al*. (1998). Biometric measures and sex were not available for birds belonging to Viterbo population.

**Figure 1.**
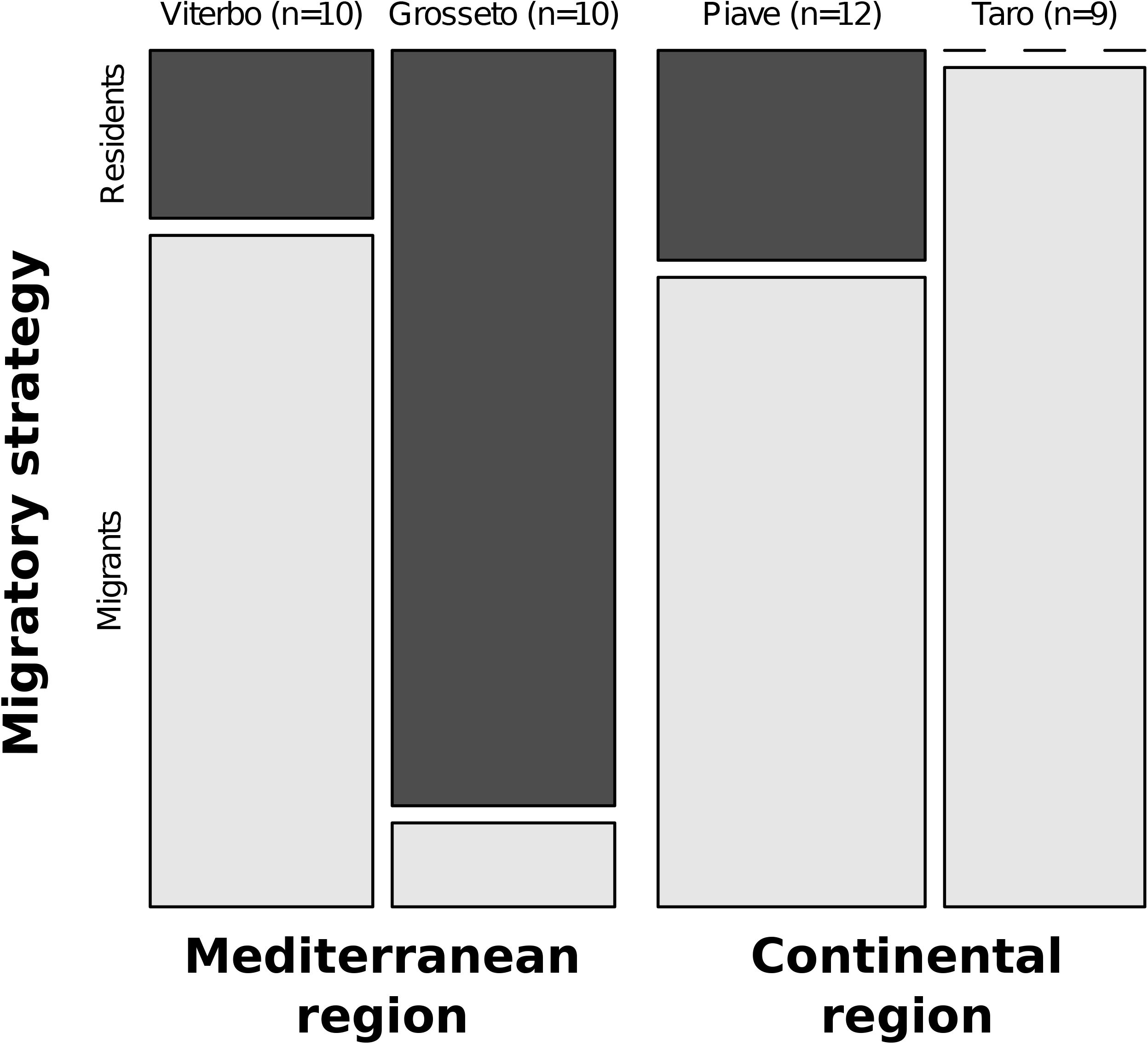
Mosaic plot of the proportion of migrant and resident Stone-curlews recorded in the four considered populations in the period 2010-2019. Data from the Taro population refer to seven animals marked with geolocators and two animals marked with GPS/UHF (see Giunchi *et al*. 2015 for details). The proportion of migrants differed significantly among populations (Firth’s bias-reduced penalized-likelihood logistic regression model: χ^2^ = 18.9, df = 3, P = 0.0003, LR test).

### Tracking data

Two types of tracking devices were used throughout the study period: geolocators and solar powered GPS tags. Geolocator data were collected in the years 2010-2011 and were already published (see Giunchi *et al*. 2015 for details). These data were taken into account solely for comparing the proportion of migrants and residents from the four populations. Two types of GPS devices (GPS-UHF and GPS-GSM) were deployed in the period 2012-2019 (see Table S1 for details). Part of the tracks considered in this study were already published in Giunchi *et al*. (2015) and in Cerritelli *et al*. (2020). The GPS were fitted on birds using a Teflon ribbon leg-loop (Rappole and Tipton, 1991) or backpack harness (Viterbo population only). The weight of the tags corresponded to the 3-4% of bird’s body weight (average bird weight = 475.4 g ± 37.3 SD, n = 23; average tag weight = 17.2 ± 2.3 g, n = 24). The weight of birds tagged in the Viterbo province was not available.

To avoid pseudo-replication (Hulbert, 1984), we randomly selected a single track for each season whenever more than two tracks were available for a given individual.

In order to minimize possible bias inherent in subjective approaches (Cerritelli *et al*. 2020), we objectively identified departure and arrival dates in both breeding and wintering areas and stopovers through the segmentation method implemented in the package ‘segclust2d’ 0.2.0 (Patin et al. 2020) in R 4.0.2 (R Core Team 2020). This method allows to find similar segments in function of the speed and tortuosity (i.e. low variability for limited periods), corresponding to stationary phases, and cluster them in a common class, corresponding to a given state. Two parameters had to be set in order to carry out the analysis (Patin et al. 2020): a minimum segment length corresponding to the minimum time that a certain behavior could be considered stationary (L_min_) and the maximum number of states (M). The optimal number of segments (K) and of states (M) are then chosen by means of model selection according to the Bayesian information criterion (BIC). Track segmentation was performed in two steps. In the first step, aimed at identifying the onset and the end of migration, we considered for each individual a tracking period ranging from 20 days before the presumed departure to 20 days after the presumed arrival, as inferred by the visualization of the tracks. The track was regularized (Patin *et al*. 2020) using ‘adehabitatLT’ 0.3.25 R-package (Calange, 2006) at 1 fix every 60 or 180 minutes to obtain the best trade-off between the maximum number of tracks available for the analyses and the minimum time lag between fixes (Table S1). The tracks were segmented by setting the L_min_ so that the minimum time of stationarity was 12 h, the maximum number of states M = 3 (residency in breeding and wintering areas and migration states).

In the second step of the segmentation process, we used the migratory tracks identified in the previous step in order to identify the stopover periods. This analysis was performed on the tracks of 14 individuals which had at least 1 fix / 90 minutes. The tracks were regularized according to the original duty cycle (Table S1) and then segmented by considering a minimum time of stationarity of 6 h with three states (rest at stopover, foraging at stopover, migration). Stopovers were not identified using the segmentation method for six birds (Table S1, individuals marked with an asterisk), as the duty cycles of their tags included switching off the instrument for 8 hours during the day. For these birds, we considered as stopovers the group of fixes collected during at least 12 h and spaced < 4 km apart, i.e. the third quartile of the distribution of the distances calculated among locations identified as stopovers for the remaining birds using the clustering-segmentation method (median = 2.1 km, first-third quartile = 0.7-4.0 km, min-max = 0-17.7 km).

After the identification of migratory and stopover periods, we calculated: the total migration distance as the sum of distances between all fixes, excluding movements in stopover areas, the straightness index, i.e. the ratio between the straight distance between wintering and breeding site and the total distance traveled by the birds (Batschelet, 1981), the number of stopovers, the time spent in stopovers, the total migration length (including the stopover period) and the time spent flying. All distance measurements were performed by means of the orthodromic Vincenty ellipsoid method using the R-package ‘geosphere’ 1.5-10 (Hijmans, 2019). Every location was classified as nocturnal (from sunset to sunrise) or diurnal considering nautical crepuscule, by means of the R-package ‘amt’ 0.1.2 (Signer *et al*. 2019) in order to verify whether this species migrates mostly during the night (Vaughan & Vaughan-Jennings, 2005).

### Regional weather data

The centroids of breeding areas for each individual were calculated and the weather data recorded in each location were extracted from the European Centre for Medium-Range Weather Forecasts European Reanalysis v5-Land data (ECMWF ERA5-Land; spatial resolution 0.1° x 0.1°; temporal resolution 1 h). As the sample size of tracked birds was relatively small, we considered a small set of variables which are known to affect the likelihood of departure (see e.g. Liechti 2006, Conklin & Battley 2011, Dänhardt & Lindström 2001, Grönroos *et al*. 2012, Duijns *et al*. 2017, Burnside *et al*. 2021) to avoid data dredging. We considered the period starting about 25 days before the departure of the first bird, i.e. from 1 October until the departure of all birds from the breeding areas. The variables included in the analysis were (Table S2): 1) the daily average of the northward component of the wind speed (10 m above the ground; Vwind, m/s), 2) the daily minimum temperature (2 m above the ground; Tmin, °C), 3) the difference between the minimum temperature of the day and that of the previous one (ΔTmin, °C) and 4) the difference between the daily average of the sea surface atmospheric pressure of the day and that of the previous one (ΔPatm, hPa). The correlation among these variables, assessed with the Spearman correlation coefficients was always well below 0.5. Photoperiod was not included in the model because its effect on migration timing is well known (Gwinner 1990, Berthold 1996, Gwinner & Helm 2003) and the aim of our analysis was to assess the effect of proximate meteorological variables in modulating the start of migration in a time window when day length is permissive for migration.

### Statistical analyses

We initially tested if the proportion of migrants and residents differed in the four populations including in the analysis both birds tagged with geolocators and those tagged with GPS. We used a Firth’s bias-reduced penalized-likelihood logistic regression model as implemented in the R-package ‘logistf’ 1.24 (Heinze *et al*. 2020), with migration strategy (presence or absence of migration) as binary dependent variable and population (four-levels factor: Taro, Piave, Grosseto, Viterbo) as independent variable. Model significance was assessed using the likelihood ratio (LR) test. We considered three planned contrasts, i.e. continental versus Mediterranean populations, Taro vs Piave population and Grosseto vs Viterbo population.

The analysis of the migratory behaviour was performed by comparing birds from the Mediterranean region (one from Grosseto population and eight from Viterbo population), with birds from the Piave (n = 9) and Taro (n = 2) populations, as representative of the Continental region. We used standard linear models (LM) with departure or arrival date (Julian day, 1 = 1st of January) as dependent variable and region as predictor to investigate the relationship between timing of migration, season and area. Data from the two migratory seasons were analysed separately. To verify if this species is a nocturnal migrant and if this behaviour changes between seasons and regions, the proportion of migratory fixes recorded during the night was compared between seasons (two-levels factor: autumn vs spring) and regions (two-levels factor: Continental region vs Mediterranean region) using a Generalized Linear Mixed Model (GLMM) with binomial error distribution and bird ID as random intercept. A linear mixed models (LMM) was used to test whether the distance traveled was affected by the migratory season, the breeding region and by their interaction and to investigate the relationship between speed of migration, season and area of origin. To take into account the effect of different duty cycles in LMM, we included a weight variable in LMM of distance and speed: a weight of 1 was assigned to the animals with a duty cycle of 60 min during migration, whereas a weight equal to 0.66 was assigned to the two animals with a duty cycle of 90 min. In all LMM models, bird ID was included as random intercept.

(G)LMM were fitted using the package ‘lme4’ 1.1-26 (Bates *et al*. 2015). Significance of fixed effects was tested by means of the likelihood ratio (LR) test. Model fit, overdispersion and collinearity were checked by means of the packages ‘performance’ 0.6.1 (Lüdecke *et al*. 2020) and ‘DHARMa’ 0.3.3.0 (Harting 2020). 95% confidence intervals were estimated using the package ‘parameters’ 0.6.1 (Lüdecke *et al*. 2020). Adjusted R^2^ was reported for every linear model. Conditional and marginal R^2^ for the linear mixed models was calculated according to Nakagawa *et al*. (2017) using the package ‘MuMIn’ 1.43.17 (Barton, 2020).

The possible effect of the considered meteorological variables on the start of migration of the Stone-curlew was investigated by means of the non-parametric Cox proportional hazard model (Moore, 2016). This model describes the probability per unit of time that an event occurs as a function of the baseline hazard and a set of covariates (fixed or time-dependent). The models were fitted using the R-package ‘survival’ 3.2-11 (Therneau 2020, Therneau & Grambsch 2000). Data were clustered by bird ID and robust variance was estimated using a jackknife approach (Therneau 2020, Therneau & Grambsch 2000). The predictors included in the two models were the breeding region (two-levels factor: Continental region vs Mediterranean region) and four time-dependent covariates (Vwind, ΔPatm, Tmin and ΔTmin; Table S2). The breeding region was included in the model to take into account the difference in migratory length among the populations belonging to the two areas (see Results). Model selection was performed according to the corrected Akaike Information Criterion (AICc; Burnham & Anderson 2002) using the ‘MuMIn’ package (Barton, 2020). We considered all combinations of the three time-dependent predictors while keeping the stratum covariate fixed. Given the relatively small available sample size, we reduced model complexity by not considering any interaction and by limiting the maximum number of terms to three (intercept included). The model with the lowest AICc value was used for inferences, provided that the null model was not included in the set of models within two units from the best one (Burnham & Anderson 2002). For each model we calculated the Akaike weight w_i_, representing the probability of the model given the data (Burnham & Anderson 2002). Model significance was tested using the Robust Score (RS) test while goodness-of-fit was evaluated by means of model concordance (Therneau 2020, Therneau & Grambsch 2000). RS test, model concordance, covariate coefficients and estimated hazard ratios of departure after 1 SD increase of the variables included in the model were reported for all models within two units from the best one (Burnham & Anderson 2002). Model assumptions were checked according to Therneau & Grambsch (2000) using the package ‘survminer’ 0.4.9 (Kassambara *et al*. 2021).

All statistical analyses were performed with R (version 4.0.2) (R Core Team 2020). All geographical analyses and plots were carried out using QGIS 3.4 (QGIS Development Team 2019).

## Results

### Migration strategy

The proportion of migrants differed significantly among populations (χ^2^ = 18.9, df = 3, P = 0.0003, Firth’s bias-reduced penalized-likelihood logistic regression model, LR test; Fig. 1). As expected, migratory birds were more common in continental than in Mediterranean populations (χ^2^ = 7.4, P = 0.007). The proportion of migrants was not different between the two continental populations (Taro vs Piave: χ^2^ = 2.1, df = 1, P = 0.1), while, contrary to the expectations, it was significantly higher in Viterbo than in Grosseto population (χ^2^ = 9.8, df = 1, P = 0.002).

### Migration routes, speed and timing

From the 20 GPS-tagged Stone-curlews identified as migrants, we reconstructed a total of 33 migratory tracks: 14 in autumn and 19 in spring. Migration turned out to be rather short in both seasons (on average less than 1000 km covered in less than five days; Table 1). Migrant Stonecurlews spent their wintering period within the Mediterranean basin (Fig. 2), i.e. in Sardinia (n = 4), Sicily (n = 3), Tunisia (n = 12) and Libya (n = 1).

**Figure 2.**
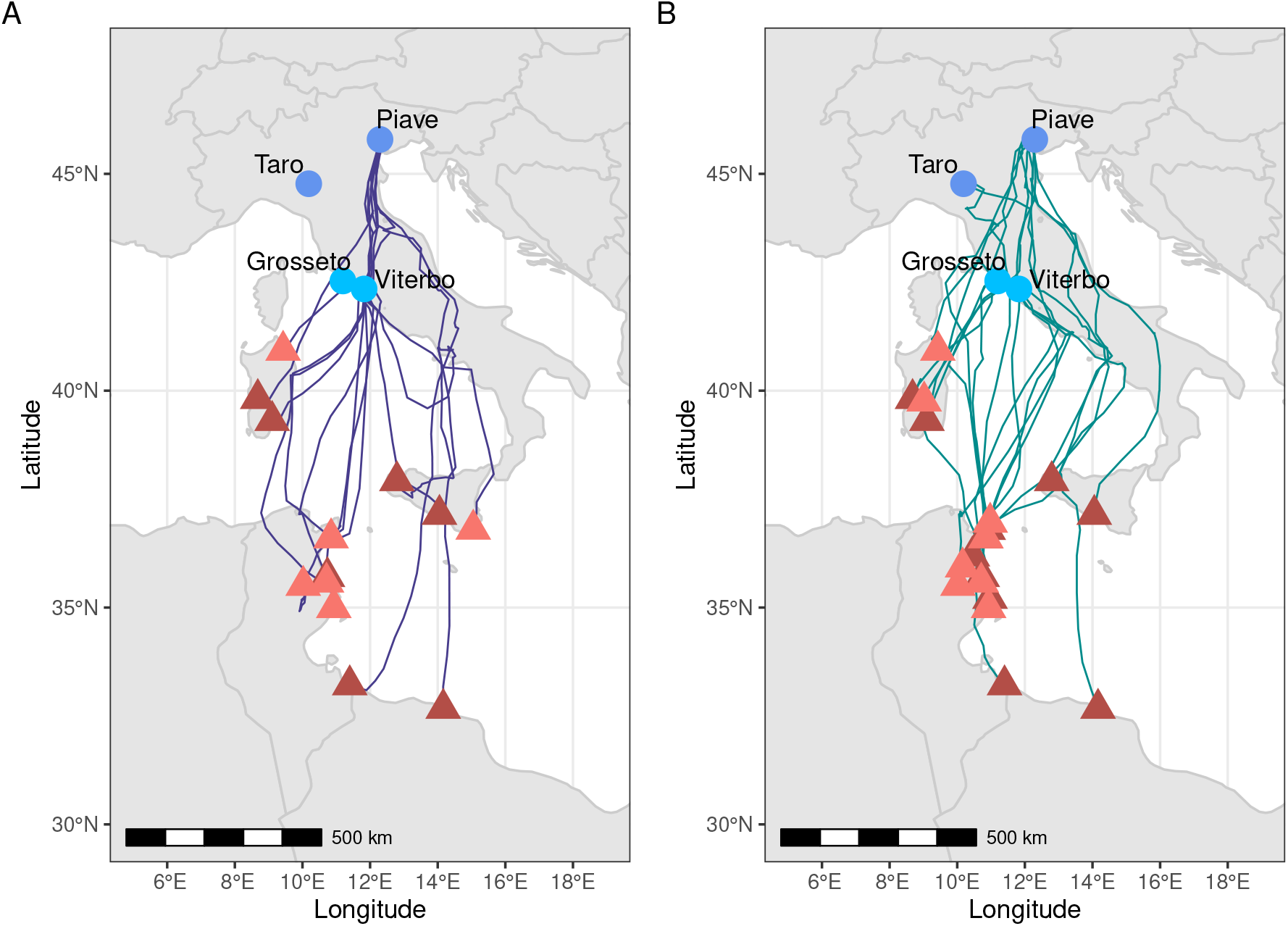
Capture areas (dots), migration tracks and overwinter locations (triangles) of Stonecurlews equipped with GPS in Italy in the period 2012-2019. **A**, autumn migration (n = 14 individuals); **B**, spring migration (n = 19 individuals). Darker symbols: continental populations; lighter symbols: Mediterranean populations.

**Table 1.** Summary statistics of individual autumn and spring migratory tracks of 20 migrant stonecurlews equipped with GPS in Italy in the period 2012-2019. Migration speed was calculated as the total distance traveled divided by the migration length, whereas the actual speed of migration was the result of the total distance traveled between successive stopovers divided by the number of traveling days.

|  | Autumn migration (*n* = 14)<br>Median (IQR)<br>Range | Spring migration (*n* = 19)<br>Median (IQR)<br>Range |
| --- | --- | --- |
| Departure date | 25 Nov (22 Nov - 3 Dec)<br>25 Oct - 31 Dec | 3 March (28 Feb - 14 Mar)<br>8 Feb - 10 Apr |
| Arrival date | 2 Dec (27 Nov - 6 Dec)<br>28 Oct - 2 Jan | 8 Mar (2 Mar - 21 Mar)<br>9 Feb - 12 Apr |
| Total duration of migration (days) | 3.3 (2.2 - 6.0)<br>1.1 - 17.5 | 4.3 (1.4 - 8.1)<br>0.5 - 12.3 |
| Time spent flying (days) | 1.4 (1.1 - 2.0)<br>0.7 - 2.7 | 1.1 (0.9 - 2.4)<br>0.5 - 3.3 |
| Time spent in stopovers (days) | 1.9 (0.9 - 3.8)<br>0.4 - 14.8 | 2.7 (0.6 - 6.2)<br>0 - 9.9 |
| Number of stopovers | 2 (2 - 3)<br>1 - 4 | 2 (1 - 5)<br>0 - 7 |
| Migration length (km) | 956.5 (798.9 - 1160.6)<br>252.2 - 1601.0 | 977.0 (794.2 - 1206.6)<br>350.4 - 1641.8 |
| Migration speed (km/day) | 279.0 (230.8 - 409.2)<br>86.7 - 525.0 | 306.9 (139.4 - 645.4)<br>38.5 - 1320.8 |
| Actual speed (km/day) | 709.2 (584.8 - 852.9)<br>360.2 - 954.2 | 736.7 (514.9 - 963.9)<br>389.4 - 1401.5 |
| Straightness index | 0.90 (0.85 - 0.93)<br>0.59 - 0.98 | 0.90 (0.83 - 0.92)<br>0.66 - 0.95 |

All birds followed a straight and direct migratory route (most straightness indexes > 0.80) both in winter and in spring (Table 1). The only exceptions to this pattern were TK2343 from the Piave population which showed a clear detour during autumn migration leading to a relatively low straightness index (0.59; Fig S1A) and GR06 from Grosseto population during spring migration which showed a moderate detour first arriving at the breeding site (0.67; Fig S1B).

The departure and arrival dates showed a high degree of variability in both migratory seasons, ranging from the last week of October to the end of December/early January in autumn, and from early February to early April in spring (Table 1). Departure and arrival dates were significantly different between the two areas both in autumn and in spring (autumn departure: F_1,12_ = 9.3, P = 0.01; autumn arrival: F_1,12_ = 8.3, P = 0.014; spring departure: F_1,17_ = 5.2, P = 0.035; spring arrival: F_1,17_ = 7.4, P = 0.015; LM, Fig. 3). In autumn, continental Stone-curlews departed on average ca. 25 days earlier than Mediterranean ones, while in spring their onset of migration was delayed by about 10 days

**Figure 3.**
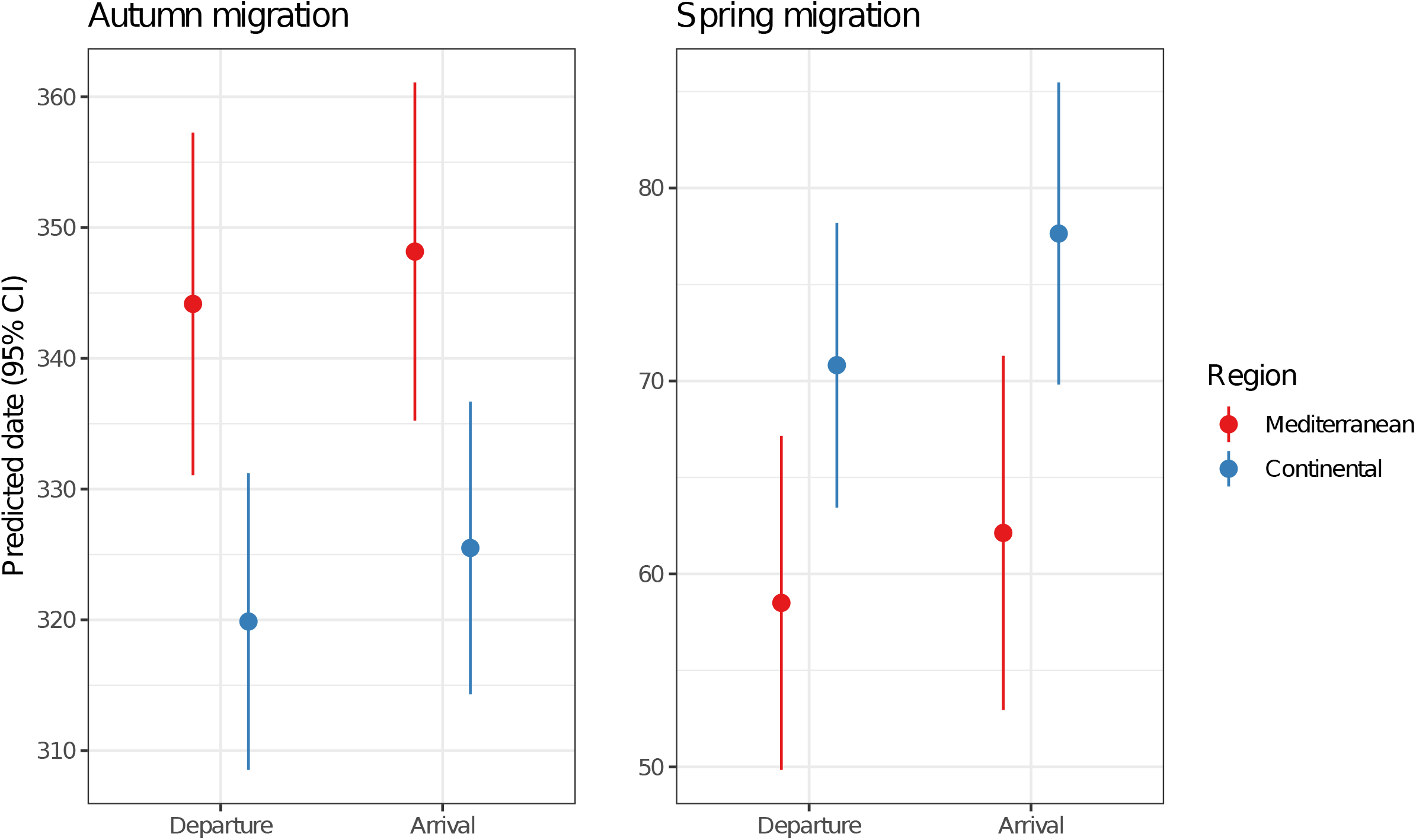
Plot of the effects of breeding area on departure and arrival of autumn and spring migration of Stone-curlews equipped with GPS in Italy in the period 2012-2019. Results from the linear models with the departure or arrival date (Julian day, 1 = 1^st^ of January) as dependent variable and region as predictor. Departure and arrival dates were significantly different between the two regions both in autumn and in spring (autumn departure: F_1,12_ = 9.3, P = 0.01, adjusted R^2^ = 0.39; autumn arrival: F_1,12_ = 8.3, P = 0.014, adjusted R^2^ = 0.36; spring departure: F_1,17_ = 5.2, P = 0.035, adjusted R^2^ = 0.19; spring arrival: F_1,17_ = 7.4, P = 0.015, adjusted R^2^ = 0.26).

Most migratory movements took place at night, as expected, and the proportion of nocturnal fixes tended to be higher in autumn (χ^2^ = 7.2, df = 1, P = 0.007; GLMM, LR test; Fig. S2) while no differences between regions were recorded (χ^2^ = 0.099, df = 1, P = 0.8).

Migration length was significantly affected by the region of origin (χ^2^ = 12.3, df = 1, P = 0.0004; LMM, LR test), with individuals from continental populations covering longer distances than the Mediterranean ones (Fig. 4), but was not affected by the migratory season (χ^2^ = 0.42, df= 1, P = 0.5) nor by the interaction between season and region (χ^2^ = 0.30, df = 1, P = 0.6).

**Figure 4.**
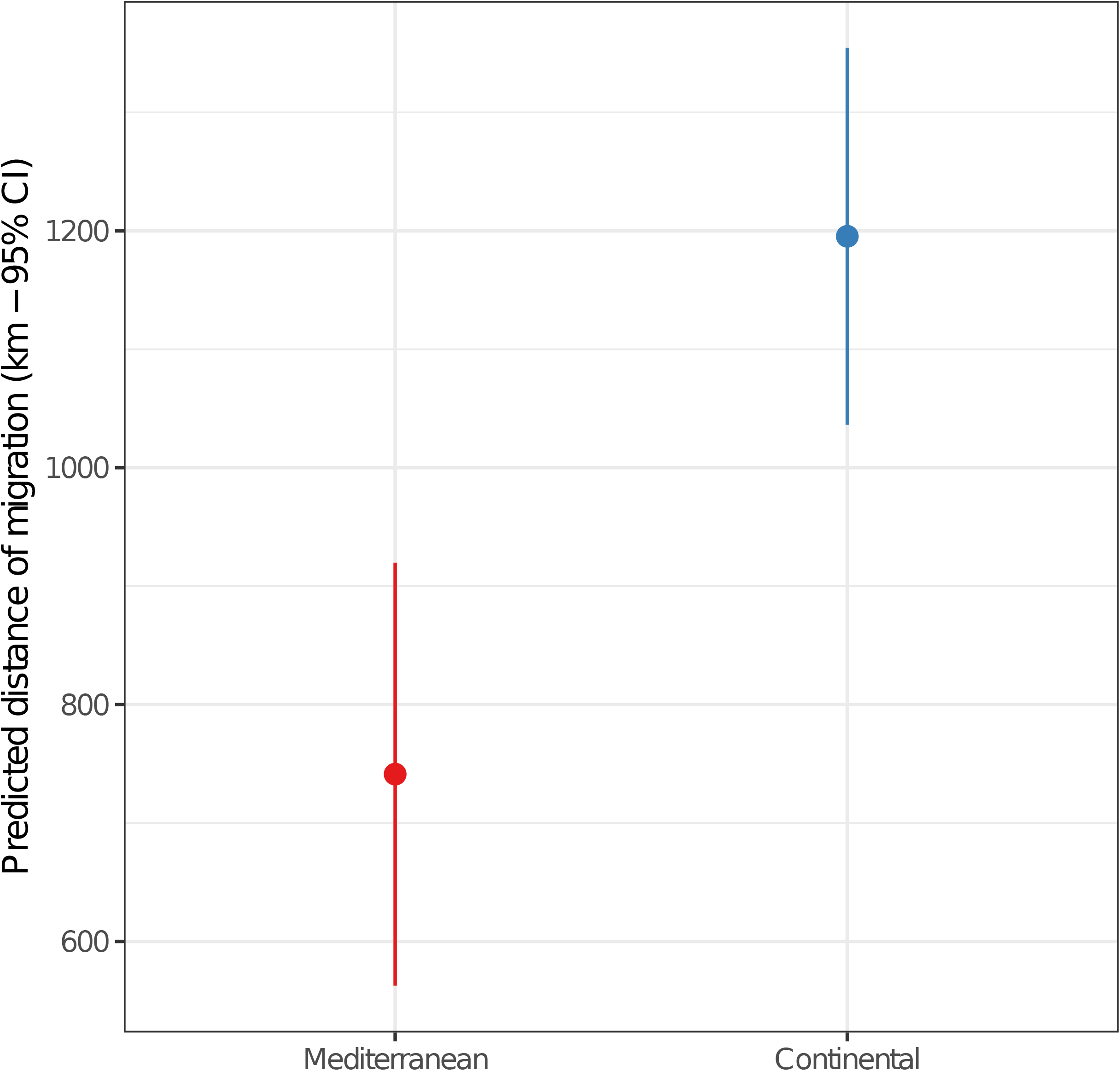
Plot of the effects of the breeding region on migration length recorded in Stone-curlews equipped with GPS in Italy in the period 2012-2019. Results from a linear mixed model with season (two-levels factor: autumn vs spring), region (two-levels factor: continental region vs Mediterranean region) and their interaction as predictors and bird ID as random intercept. Migration length was significantly different between regions (χ^2^ = 12.3, df = 1, P = 0.0004, LR test), but was not affected by migratory season (χ^2^ = 0.42, df= 1, P = 0.5) nor by the interaction between season and region (χ^2^ = 0.30, df = 1, P = 0.6). SD of the random intercept = 239.9, conditional R^2^ = 0.88, marginal R^2^ = 0.42, number of observations = 33, number of birds = 20.

The total duration of migration was rather short, often less than five days, and the median duration of active flight was about one day only (Table 1). The number of stopovers was rather variable in both seasons: usually it did not exceed two (Table 1), but two birds (VT42A, VT873) did not stop during spring migration and one bird (T25) stopped seven times during the same migration. Median stopover length was < 3 days, even though in some cases birds could stop for more than one week. The median of the total migratory speed was 300 km/day but active flight migration segments were often covered at more than 700 km/day (Table 1).

The speed of migration was significantly affected by the interaction between breeding region and migration season (χ^2^ = 5.1, df = 1, P = 0.02; LMM, LR test). Individuals from continental populations did not change their speed between seasons, whereas Mediterranean ones tended to migrate faster in spring (Fig. 5).

**Figure 5.**
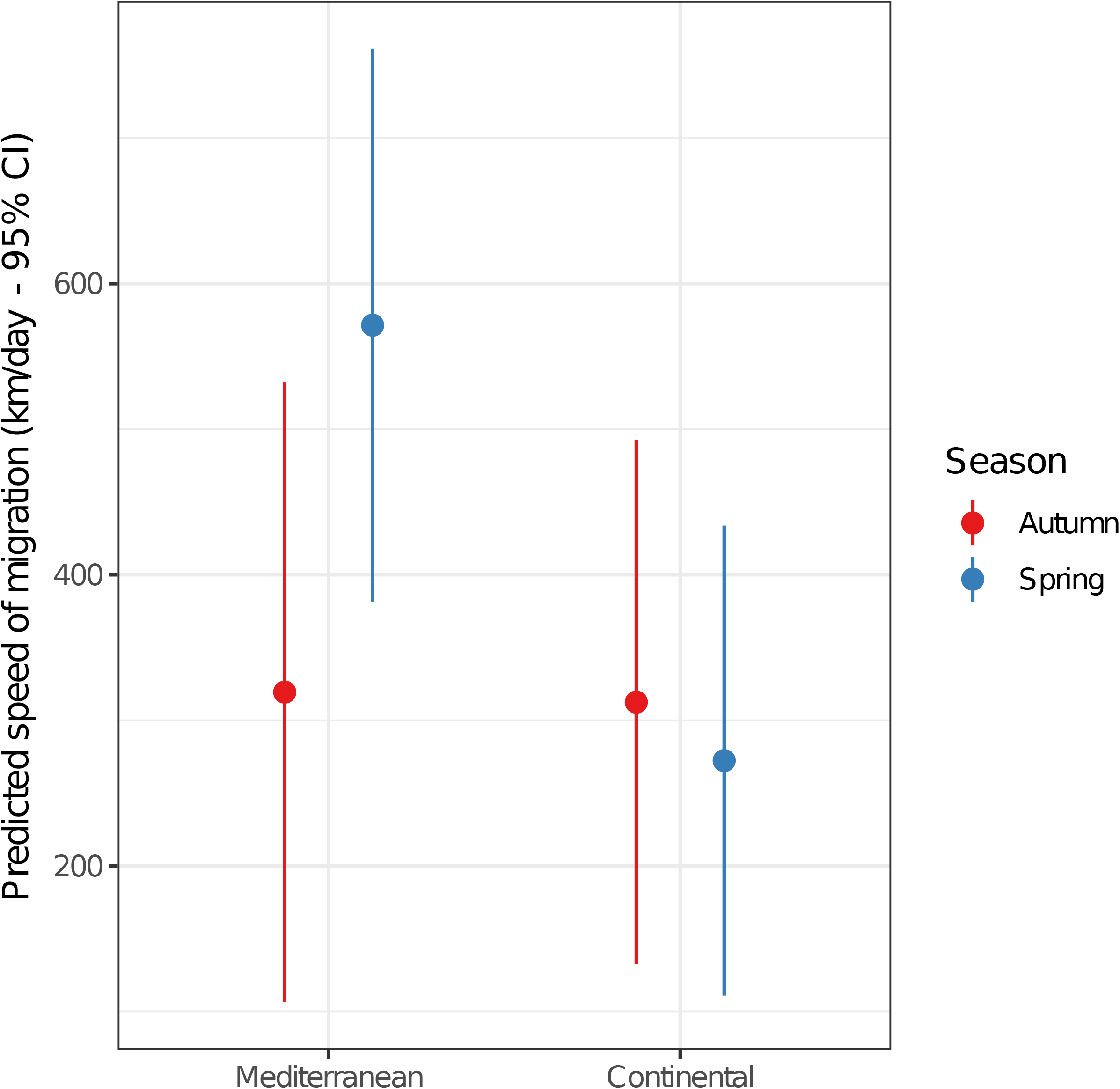
Plot of the effects of breeding region and migratory season on the speed of migration recorded in Stone-curlews equipped with GPS in Italy in the period 2012-2019. Results from a linear mixed model with season (two-levels factor: autumn vs spring), region (two-levels factor: continental region vs Mediterranean region) and their interaction as predictors and bird ID as random intercept. The effect of the interaction between season and region was significant (χ^2^ = 5.1, df = 1, P = 0.02, LR test). SD of the random intercept = 201.6, conditional R^2^ = 0.66, marginal R^2^ = 0.18, number of observations = 33, number of birds = 20.

### Effects of meteorological variables on the start of autumn migration

The analysis on the effects of meteorological factors on the start of autumn migration was performed on 14 birds (eight from the Continental region and six from the Mediterranean region). The most supported model among the considered set included the daily minimum temperature (Tmin) and the daily averaged northward wind component (Vwind) (Table 2). The effect of temperature seems to be particularly strong, as this is the only covariate included in the second model still within two units from the best one (Table 2). Both models were highly significant and have a high coefficient of concordance. As expected, the likelihood of departure increased when both Tmin and Vwind decreased, i.e. birds tended to depart in cold days when wind blew in a southerly direction (Fig. 6).

**Figure 6.**
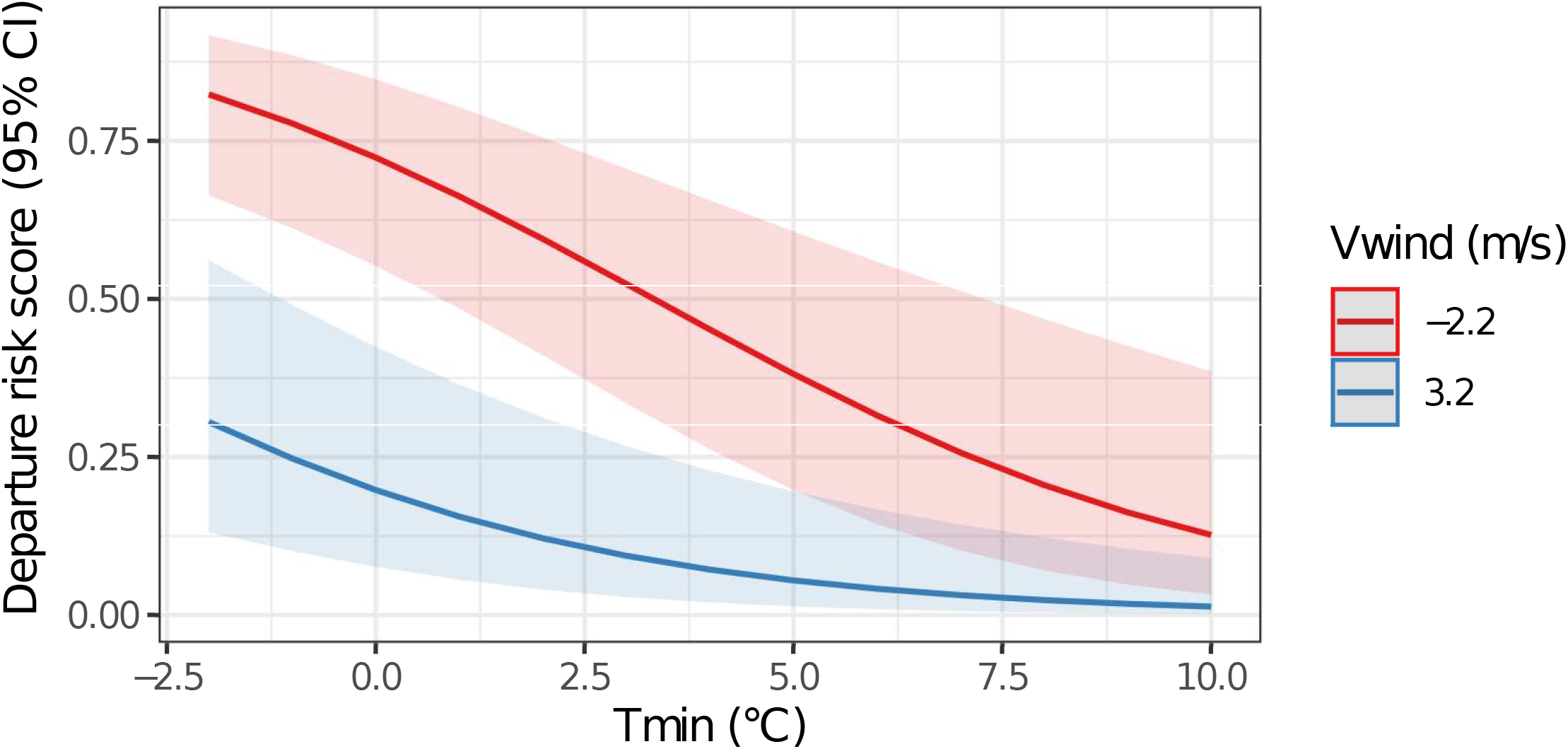
Plot of the effects of daily minimum temperature (Tmin) and daily averaged northward wind component (Vwind) on the departure risk score for autumn migration of Stone-curlews equipped with GPS in Italy in the period 2012-2019. Results from the best Cox proportional hazard models (see Table 2 for further details). Number of birds = 14.

**Table 2.**
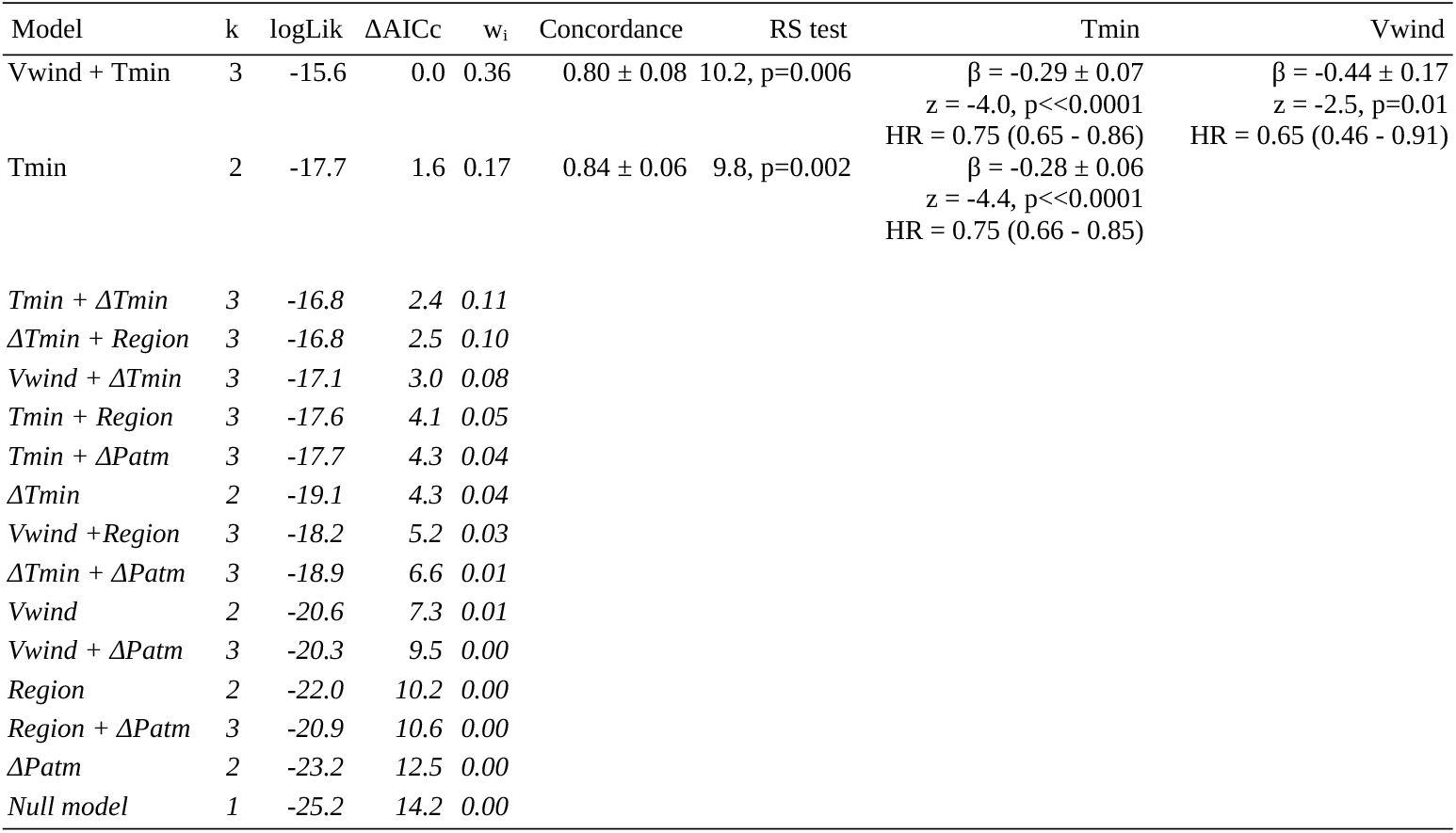
Comparison of non-parametric Cox proportional hazard models developed to identify the meteorological variables affecting the likelihood of starting autumn migratory movements by Stone-curlews equipped with GPS in Italy in the period 2012-2019. Concordance (± SE), model significance [Robust Score (RS) test], model coefficients (β ± SE) and relative significance, estimated hazard ratio (HR) of departure after 1 SD increase of the variables included in the model (95% coefficient intervals) are reported only for models within two units from the best model. Predictors included in the models: Vwind, daily averaged northward wind component (m/s); Tmin, daily minimum temperature (°C); ΔTmin, difference between the minimum temperature of the day and that of the previous one (°C); ΔPatm, difference between the average sea level atmospheric pressure of the day and that of the previous one (hPa). k, number of parameters; logLik, loglikelihood; ΔAICc, difference in AICc between a given model and the model with the lowest AICc; w_i_, Akaike weights. Number of observations = 802; Number of birds = 14.

## Discussion

This study reports one of the few detailed investigations of the migratory behaviour of a Mediterranean migrant and the first of this kind for the Stone-curlew. The results highlight the significant intra- and inter-population variability of the migratory behaviour of Italian breeding populations, which is not fully explained by breeding latitude and suggest an important role of temperature and wind conditions in modulating the timing of autumn migration.

### Migration strategy

Three of the four considered populations were found to be partially migratory (Chapman *et al*. 2011). This result is not surprising as partial migration is a fairly common strategy among shortdistance migrants (Pulido *et al*. 1996, Newton 2008). The analysis of the proportion of migrating individuals showed significant interpopulation variability, as recorded for other migratory species in the Mediterranean basin (de la Morena *et al*. 2015, Monti *et al*. 2018). Individuals belonging to northern populations (Taro and Piave) were more likely to undertake migration than populations belonging to the centre of the peninsula, in agreement with our expectations based on data from other Palaearctic migrants (Newton & Dale 1996, Newton 2008) and with what has been previously suggested for the Stone-curlew (Brichetti & Fracasso 2004, Biondi & Pietrelli 2015). The proportion of migrants was not different between the Piave and Taro populations, but the Taro population was the only one in which all tagged birds were found to be migrants. The relatively small number of birds tagged in the Taro could at least partly explain these results, as resident birds were probably rare, while an effect of the low accuracy of the geolocators used in the Taro population seems unlikely (see Giunchi *et al*. 2015 for details). Since individuals from the Taro population were tagged earlier than those from the Piave population, however, we cannot exclude that the small difference in the proportion of residents in the two populations could be interpreted as an ongoing change in the migratory strategies of the northern populations of the species, as recently observed for other birds migrating over short distances (Doswald *et al*. 2009, Pautasso 2012, Ambrosini *et al*. 2016, Tellería *et al*. 2016, Podhrázský *et al*. 2017).

The difference in the proportion of migrants between the two southernmost populations (Viterbo and Grosseto) was on the other hand unexpected,, as their respective breeding areas are very close to each other (about 57 km) and subject to comparable climatic conditions. We could speculate that this result could be due to intrinsic differences between tagged individuals possibly related to their sex and age, which are known to influence the migration strategy of birds (Gauthreaux 1982, Terrill & Able 1988, Newton 2008, Chapmann *et al*. 2011, Hegemann *et al*. 2019). We did not record the sex of Viterbo birds, but data collected on the remaining populations did not show a consistently different pattern in the migration strategy of males and females (data not shown). Furthermore, all birds considered in this study were adults (> one year old). Interestingly, the only exception was the single migrant in the Grosseto population, which migrated when it was one year old and then became resident for the next two years of monitoring. This suggests that age and/or competitive ability may indeed play a role in modulating the migratory behaviour of Stone-curlews, as observed in other partial migrants (see Newton 2008, Chapmann *et al*. 2011, Hegemann *et al*. 2019 for references), but it is not very helpful in understanding the unexpected migratory propensity of Viterbo birds. It can not be excluded that this result is due to genetic differences between the two populations, even though previous genetic analyses (not including the Viterbo population) suggested that the gene flow between Italian Stone-curlew populations is significant (Mori *et al*. 2017). Alternatively, it could be hypothesised that the resources available in the Viterbo area during the non-breeding season are more limited than in the Grosseto area, e.g. due to a more widespread use of intensive agricultural practices, thus leading a higher fraction of individuals to migrate (Cox 1985, Boyle 2008, Jahn *et al*. 2010). Beyond the possible explanations, this result highlights a high local variability in the migration strategy of this species that deserves further investigation.

### Migration routes, speed and timing

Most birds followed a relatively straight and direct route both in autumn and in spring. Migratory movements were limited within the Mediterranean basin, but wintering sites showed a wide latitudinal range (from Sardinia to Libya) compared to the latitudinal range of breeding areas and to the scale of migratory movements. The large overlap of wintering sites between northern and central populations indicates that the migration pattern is telescopic and leads populations that are allopatric during the breeding period to overwinter in sympatry (Netwon 2008, Borras *et al*. 2011, Arizaga & Tamayo 2013). This pattern suggests a weak connection between breeding and non-breeding areas (migratory connectivity, Webster *et al*. 2002) which can have significant implications for the conservation of the species (Cresswell 2014, Finch *et al*. 2017). In this regard, it is interesting to observe that all tagged migrant Stone-curlews spent the winter season in areas that probably already hosted resident populations of the species (Delany *et al*. 2009, Brichetti & Fracasso 2004, Tinarelli *et al*. 2009, Hume & Kirwan 2020).

Our data were only partially in agreement with the expectation that total migration speed should be higher in spring than in autumn (Kokko 1999, Nilsson *et al*. 2013, Zhao *et al*. 2017, Schmaljohann 2018). Spring migration was indeed faster only for Mediterranean Stone-curlews, whereas no difference was observed for continental birds. Population variability in the difference in overall migration speed between seasons has been observed in other species (see e.g. Vansteelant *et al*. 2015, Schmaljohann 2018). The variability recorded in this study was relatively small, so we might assume that Stone-curlews did not significantly change their migration strategies in different seasons, given the short migration distance. Some data collected on shorebirds of different sizes also seemed to suggest that the seasonal difference in migration speed tended to be smaller for larger species (Zhao *et al*. 2017). It cannot be ruled out, however, that Stone-curlews do indeed adopt different strategies in spring and autumn, but no difference in migratory speed could be detected for continental birds due to the effect of unfavourable weather conditions encountered en route that may have delayed or interrupted the migratory journey (see e.g. Newton 2008, Shamoun-Baranes *et al*. 2010, Shamoun-Baranes *et al*. 2017). This effect might have been less relevant for Mediterranean individuals, because the greater proximity between the wintering and breeding areas might allow them to better adjust their timing of migration in function of weather conditions they expect to find en route and in the area of arrival (Lehikoinen *et al*, 2004, Rubolini *et al*. 2007, Møller *et al*. 2008, Knudsen *et al*. 2011). In this regard, it is interesting to note that birds from both areas tended to migrate during the day more frequently in spring than in autumn, as indicated by the significantly lower fraction of migratory fixes recorded during the night in spring. This extension of nocturnal migration into the daytime, which has been recorded for other nocturnal migrants especially when crossing ecological barriers (Newton 2008, Adamik *et al*. 2016, Malminga *et al*. 2021), might suggest a tighter migratory schedule in spring and therefore seems to be in agreement with a time-minimization strategy.

Even though the latitudinal range of migratory movements was quite small, we observed a significant effect of breeding latitude on the timing of migration. Continental birds departed significantly earlier than Mediterranean ones in autumn while the opposite occurred for spring migration and this was expected according to the available data on a wide range of migratory species (e.g. Newton 2008, Conklin *et al*. 2010, Linek *et al*. 2021). The pattern observed for arrival dates was the same as that observed for departure dates probably because the difference in the migration distance between the two areas was quite small. Interestingly, all autumn departures occurred after the end of primary moult estimated using data from the Taro population (17^th^ October ± 14 days SD, Giunchi *et al*. 2008). This indicates that at southern latitudes animals have sufficient time to complete the moulting of the remiges before migrating and do not need to suspend the moult as appears to be the case at northern latitudes (see Giunchi *et al*. 2008 for details).

The timing of migration was quite variable as both autumn and spring departures occurred over a period of two months. This variability is in line with the observed flexibility of the migratory schedule of short-distance migrants (Pulido & Widmer 2005, Newton 2008, Pulido & Berthold 2010, Monti *et al*. 2018a) which are often characterized by a sort of “relaxed migratory systems” (see Monti *et al*. 2018a). Contrary to the expectations (Shamoun-Baranes *et al*. 2006, Nilsson *et al*. 2013) and to what has been observed in other shorebird species (e.g. *Vanellus gregarius*, Donald *et al*. 2021), the variability of departure dates was comparable between autumn and spring. The wide interval of departures observed in both seasons could be at least partially due to a different timing of migration between age or sex classes (differential migration, Ketterson & Nolan 1993, Newton 2008). We can not exclude an effect of sex, as the sex of some of the tracked birds was not available while the effect of age seems rather unlikely as all tracked birds were adults (> 1-year old).

### Effects of meteorological variables on the start of autumn migration

The analysis of the meteorological proximate factors affecting the timing of migration suggests that departure dates are affected in particular by temperature and wind conditions. As expected, birds tended to depart in tailwind conditions (Åkesson & Hedenström 2000, Liechti 2006, Gill *et al*. 2014, but see Eikenaar & Schmaljohann 2015, Schwemmer *et al*. 2021), but the greatest effect on the probability to start autumn migration is due to the daily minimum temperature, probably for its negative effect on the availability of invertebrates (Eggleton *et al*. 2009, Abram *et al*., 2017) and on the cost of thermoregulation for a species with a low resting metabolic rate (Duriez *et al*. 2010). The effect of temperature on migratory departure has been documented in several long- and short distance migrants (e.g. Xu & Si 2019, Klinner & Schmaljohann 2020, Liek *et al*. 2021). In a medium-distance steppic migrant, the Asian houbara *Chlamydotis macqueenii*, the effect of temperature was stronger in spring rather than in autumn, as departure decisions were less repeatable for temperature in autumn rather than in spring (Burnside *et al*. 2021). Unfortunately, this pattern cannot be verified in our species, as the small sample size available and the latitudinal spread of wintering areas did not allow robust modelling of the effect of temperature on the onset of spring migration. In addition, the lack of a significant number of repeated migration tracks did not allow an estimation of the repeatability of individual migration timing.

The relatively large increase in the probability of departure when temperatures were near or below 0 °C might suggest that Stone-curlews may tend to delay departure until winter conditions worsen abruptly (Haila *et al*. 1986, Netwon 2008). This strategy could be effective at the considered latitudes, because the relatively mild environmental conditions combined with the short migration distance could allow for a relatively relaxed timing of the start of migration. This seems to be supported by the wide time interval of autumn departures, some of which occurred even in the second half of December. According to this hypothesis, it can be predicted that the frequency of migratory birds in the considered populations will decrease in the near future due to the effect of global warming in the Mediterranean (Jenni & Kéry 2003, Gordo 2007). This could consequently modify the present distribution of the species throughout the year and should be taken into account when targeting conservation measures, as the ecological needs of the Stone-curlew could change between breeding and non-breeding seasons.

## Acknowledgements

This research was partially funded by The Taro River Regional Park, Gruppo Ornitologico Maremmano, Ekoclub Treviso, Department of Biology (University of Pisa). We would like to thank all the people who helped us in the field, and in particular: P. Berra, C. Caccamo, E. Pollonara, L. Passalacqua, F. Rodriguez-Godoy. All protocols performed in studies involving animals comply with the ethical standards and Italian laws on animal welfare. All procedures involving animals were approved by the Italian Istituto Superiore per la Protezione e la Ricerca Ambientale (ISPRA).

