## Supplementary material for "Eurasian Stone-curlews Burhinus oedicnemus breeding in Italy show a remarkable inter and intra-population variability of migratory behaviour"

**Electronic Supplementary Material**

**Table S1.** Summary information of Stone-curlews equipped with GPS in Italy in the period 2012-2019. The intervals between consecutive fixes are reported only for migrant individuals. The tags of the six birds marked with an asterisk were switched off for eight hours during the day. Track regularization was performed for all birds before the track segmentation aimed at identifying the onset and the end of migration (see the main text for further details)

| ID | Population | Region | Year of capture | Migratory strategy | GPS tag manufacturer | Interval between consecutive fixes (min) | Interval between consecutive fixes after track regularization (min) | Tracking duration (days) |
| --- | --- | --- | --- | --- | --- | --- | --- | --- |
| GR6 | Grosseto | Mediterranean | 2014 | Migrant | Ecotone | 15, 30 | 60 | 1135 |
| P01 | Piave | Continental | 2015 | Migrant | Ecotone | 30 | 60 | 383 |
| P04 | Piave | Continental | 2014 | Migrant | Ecotone | 15 | 60 | 564 |
| P06 | Piave | Continental | 2015 | Migrant | Ecotone | 30 | 60 | 818 |
| P07 | Piave | Continental | 2014 | Migrant | Ecotone | 15, 30, 60 | 60 | 300 |
| P20 | Piave | Continental | 2015 | Migrant | Ecotone | 30 | 60 | 802 |
| TK2314 | Piave | Continental | 2019 | Migrant | Ecotone | 15, 30, 60 | 60 | 577 |
| TK2343 | Piave | Continental | 2019 | Migrant | Ecotone | 15, 30, 60 | 60 | 717 |
| TK2347 | Piave | Continental | 2019 | Migrant | Ecotone | 15, 30, 60 | 60 | 494 |
| TK2349 | Piave | Continental | 2019 | Migrant | Ecotone | 15, 30, 60 | 60 | 533 |
| T24 | Taro | Continental | 2012 | Migrant | Ecotone | 30 | 60 | 233 |
| T25 | Taro | Continental | 2012 | Migrant | Ecotone | 30 | 60 | 170 |
| VT_2E7 | Viterbo | Mediterranean | 2019 | Migrant | TechnoSmart | 60* | 180 | - |
| VT_42A | Viterbo | Mediterranean | 2018 | Migrant | TechnoSmart | 60* | 180 | 1275(ongoing) |
| VT_4DD | Viterbo | Mediterranean | 2019 | Migrant | TechnoSmart | 60* | 180 | 605 |
| VT_873 | Viterbo | Mediterranean | 2018 | Migrant | TechnoSmart | 60* | 180 | 1212 |
| VT_907 | Viterbo | Mediterranean | 2019 | Migrant | TechnoSmart | 60* | 180 | 573 |
| VT_A8A | Viterbo | Mediterranean | 2018 | Migrant | TechnoSmart | 60* | 180 | 940 |
| VT_CCB | Viterbo | Mediterranean | 2017 | Migrant | TechnoSmart | 90 | 180 | 548 |
| VT_D20 | Viterbo | Mediterranean | 2017 | Migrant | TechnoSmart | 90 | 180 | 1275 |
| GR01 | Grosseto | Mediterranean | 2015 | Resident | Ecotone | - | - | 701 |
| GR02 | Grosseto | Mediterranean | 2014 | Resident | Ecotone | - | - | 733 |
| GR03 | Grosseto | Mediterranean | 2016 | Resident | Ecotone | - | - | 353 |
| GR04 | Grosseto | Mediterranean | 2016 | Resident | Ecotone | - | - | 321 |
| GR07 | Grosseto | Mediterranean | 2013 | Resident | Ecotone | - | - | 244 |
| GR08 | Grosseto | Mediterranean | 2013 | Resident | Ecotone | - | - | 659 |
| GR09 | Grosseto | Mediterranean | 2013 | Resident | Ecotone | - | - | 167 |
| GR10 | Grosseto | Mediterranean | 2014 | Resident | Ecotone | - | - | 944 |
| GR11 | Grosseto | Mediterranean | 2015 | Resident | Ecotone | - | - | 688 |
| P05 | Piave | Continental | 2015 | Resident | Ecotone | - | - | 639 |
| TK2345 | Piave | Continental | 2019 | Resident | Ecotone | - | - | 406 |
| TK2346 | Piave | Continental | 2019 | Resident | Ecotone | - | - | 778 |
| VT_290 | Viterbo | Mediterranean | 2017 | Resident | TechnoSmart |  | - | 240 |
| VT_96A | Viterbo | Mediterranean | 2018 | Resident | TechnoSmart |  | - | 397 |

**Table S2.** Summary statistics of the environmental variables considered in the analysis of the factors affecting the onset of autumn migration.

|  |  | Continental region (n= 375) | | | | Mediterranean region (n= 427) | | | |
| --- | --- | --- | --- | --- | --- | --- | --- | --- | --- |
| Variable | Description | Mean | Median | IQR | Range | Mean | Median | IQR | Range |
| Vwind | Daily averaged northward wind component (m/s) | -2.8 | -0.5 | -0.9 - 0.2 | -2.8 - 1.1 | -8.3 | 0.7 | -1.1 - 2.4 | -8.3 - 9.4 |
| Tmin | Daily minimum temperature (°C) | 9.4 | 9.6 | 6.6 - 12.4 | -1.8 - 17.6 | 12.6 | 13.0 | 10.4 - 14.7 | 1.7 - 21.3 |
| ΔTmin | Difference between the minimum temperature of the day and that of the previous one (°C) | -0.3 | -0.3 | -1.5 - 0.9 | -8.7 - 6.3 | -0.2 | -0.2 | -1.3 - 0.7 | -5.8 - 6.0 |
| ΔPatm | Difference between the average sea level atmospheric pressure of the day and that of the previous one (hPa) | 0.0 | 0.0 | - 0.6 - 0.6 | -4.0 - 3.5 | -0.0 | 0.0 | - 0.5 - 0.6 | -3.9 - 4.0 |

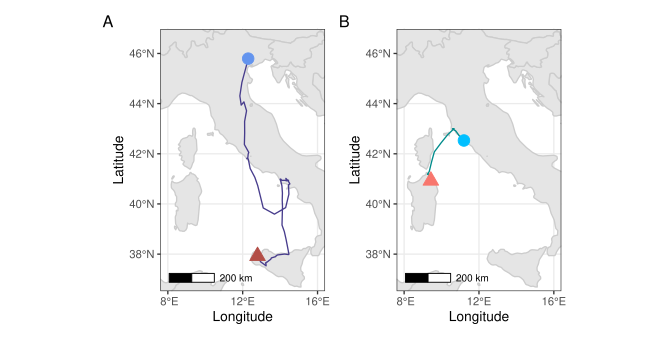

**Figure S1.** Migratory tracks of GPS-tagged Stone-curlews with a relatively low straightness index **A**, autumn migration of TK2343 (Piave population, straightness index = 0.59); **B**, spring migration of GR06 (Grosseto population, straightness index = 0.67). Dots, capture areas; triangles, overwinter locations.

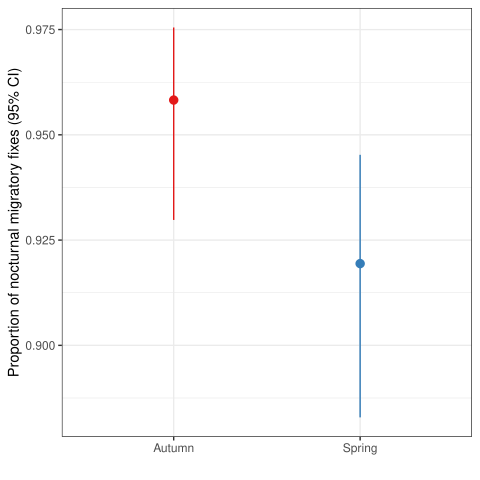

**Figure S2.** Plot of the effects of migratory season on the proportion of migratory fixes recorded during the night in Stone-curlews equipped with GPS in Italy in the period 2012-2019. Results from a generalized linear mixed model with binomial error distribution, with season (two-levels factor: autumn vs spring), region (two-levels factor: continental region vs Mediterranean region) and their interaction as predictors and bird ID as random intercept. The effect of season was significant (χ^2^ = 7.2, df = 1, P = 0.007, LR test), while the effect of region was not significant (χ^2^ = 0.099, df = 1, P = 0.8). SD of the random intercept = 0.57, conditional R^2^ = 0.12, marginal R^2^ = 0.033, number of observations = 1062, number of birds = 20.
